# Length Distribution of Ancestral Tracks under a General Admixture Model and Its Applications in Population History Inference

**DOI:** 10.1101/023390

**Authors:** Xumin Ni, Xiong Yang, Wei Guo, Kai Yuan, Ying Zhou, Zhiming Ma, Shuhua Xu

**Affiliations:** Department of Mathematics, School of Science, Beijing Jiaotong University, Beijing 100044, China; Chinese Academy of Sciences (CAS) Key Laboratory of Computational Biology, Max Planck Independent Research Group on Population Genomics, CAS-MPG Partner Institute for Computational Biology (PICB), Shanghai Institutes for Biological Sciences, CAS, Shanghai 200031, China; Institute of Applied Mathematics, Academy of Mathematics and Systems Science, Chinese Academy of Sciences, Beijing 100190, China; School of Life Science and Technology, ShanghaiTec University, Shanghai 200031, China; Collaborative Innovation Center of Genetics and Development, Shanghai 200438, China

**Keywords:** Genetic admixture, Ancestral tracks, Population history, SNP

## Abstract

As a chromosome is sliced into pieces by recombination after entering an admixed population, ancestral tracks of chromosomes are shortened with the pasting of generations. The length distribution of ancestral tracks reflects information of recombination and thus can be used to infer the histories of admixed populations. Previous studies have shown that inference based on ancestral tracks is powerful in recovering the histories of admixed populations. However, population histories are always complex, and previous studies only deduced the length distribution of ancestral tracks under very simple admixture models. The deduction of length distribution of ancestral tracks under a more general model will greatly elevate the power in inferring population histories. Here we first deduced the length distribution of ancestral tracks under a general model in an admixed population, and proposed general principles in parameter estimation and model selection with the length distribution. Next, we focused on studying the length distribution of ancestral tracks and its applications under three typical admixture models, which were all special cases of our general model. Extensive simulations showed that the length distribution of ancestral tracks was well predicted by our theoretical models. We further developed a new method based on the length distribution of ancestral tracks and good performance was observed when it was applied in inferring population histories under the three typical models. Notably, our method was insensitive to demographic history, sample size and threshold to discard short tracks. Finally, we applied our method in African Americans and Mexicans from the HapMap dataset, and several South Asian populations from the Human Genome Diversity Project dataset. The results showed that the histories of African Americans and Mexicans matched the historical records well, and the population admixture history of South Asians was very complex and could be traced back to around 100 generations ago.

## INTRODUCTION

Population admixture is a common phenomenon in human populations when previously isolated populations start to contact and interact with each other, accompanied by population migration, rising and falling of empires, trading of goods and services, and so on (Hellenthal *et al.* 2014). The history of population admixture does not fade with time, but leaves a great deal of information in the genomes of individuals from admixed populations. Population history in admixed populations thus can be recovered by utilizing the information in the genome, such as break points of recombination (Xu *et al.* 2008), admixture linkage disequilibrium (Ald) (Patterson *et al.* 2012; Loh *et al.* 2013; Pickrell *et al.* 2014) and the length of ancestral tracks (ancestral tracks) (Pool and Nielsen 2009; Pugach *et al.* 2011; Gravel 2012; Jin *et al.* 2012; Hellenthal *et al.* 2014; Jin *et al.* 2014).

The information of ancestral tracks was first used by Pool and Nielsen (which they called migration tracts) to infer the history of populations, and they also deduced the distribution of ancestral tracks under a hybrid isolation (HI, or a one pulse admixture) model with a small migration rate (Pool and Nielsen 2009). A subsequent study of Pugach *et al.* performed wavelet transform on the ancestral tracks in admixed populations to obtain the dominant frequency of ancestral tracks and compared it to those obtained from extensive simulations, to estimate the admixture time, still under the simple HI model (Pugach *et al.* 2011). Then, the study of Jin *et al.* explored admixture dynamics by comparing the empirical and simulated distribution of ancestral tracks under three typical two-way admixture models; i.e. HI model, gradual admixture (GA) model, and continuous gene flow (CGF) model (Jin *et al.* 2012). Jin *et al.* later deduced the theoretical distributions of ancestral tracks under HI and GA models (Jin *et al.* 2014). Gravel extended these studies to multiple ancestral populations and discrete migrations. However, he failed to explicitly deduce the theoretical distribution of ancestral tracks under a general situation (Gravel 2012).

Here we proposed a general model that can cover all the scenarios of an admixed population with an arbitrary number of ancestral populations and (or) arbitrary number of admixture events. In this study, we first described the general admixture model and deduced a general formula for the theoretical distribution of ancestral tracks. With this distribution, we can use maximum likelihood estimation (MLE) to estimate model parameters, and the Akaike information criterion (AIC) (Akaike 1998) or the likelihood ratio test (LRT) (Wilks 1938) to select an optimal model from candidates for the given data. We next demonstrated that the three aforementioned admixture models, namely HI, GA and CGF models in previous studies (Jin *et al.* 2012), are all special cases under our general model. Then, under these three models, we developed a method called *AdmixInfer* to estimate the admixture proportion and admixture time, and simultaneously selected the optimal model according to the principles of AIC. We further conducted extensive simulation studies to demonstrate the accuracy of the theoretical distribution of ancestral tracks under the general model, and the effectiveness of our method to estimate the parameters and select an optimal model. Finally we applied our method to African Americans and Mexicans from the HapMap phase III dataset (International Hapmap *et al.* 2010), and several South Asian populations from the Human Genome Diversity Project (HGDP) (Li *et al.* 2008) dataset.

## GENERAL MODEL

### Length Distribution of Ancestral Tracks

In our general model, population admixture is accomplished by receiving gene flow(s) from ancestral populations either continuously or discontinuously. We model this process generation by generation, in which, if the admixed population does not receive further gene flow(s) in a particular generation, we set the strength of gene flow(s) to 0. Specifically, given an admixed population started *T* generations ago, with *K* ancestral populations, let *m*_*i*_(*t*) denotes the ancestry proportion from the *ith* ancestral population that entered at the *t* generation (Figure 1), then the general model is only determined by a *K* × *T* matrix *M* = {*m*_*i*_ (*t*)}_1≤*i*≤*k*, 1≤*t*≤*T*_, which satisfies three properties: (a) *m*_*i*_(*t*) ≥ 0; (b) 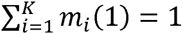 and (c)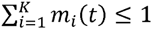, *if* 2 ≤ *t* ≤ *T*.

**Figure 1.**
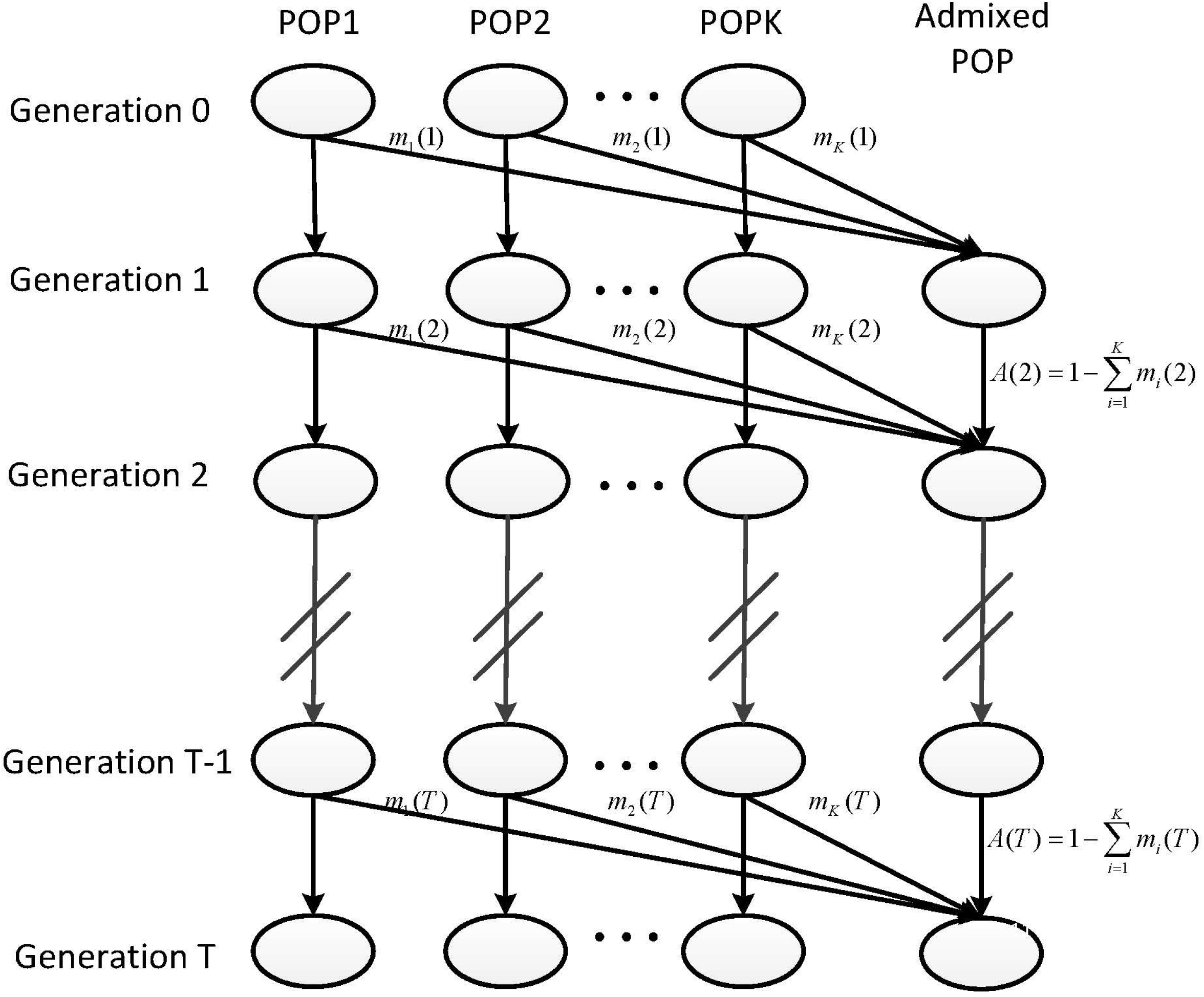
The general admixture model. Here we illustrated an admixed population with *K* ancestral populations, which started to admix *T* generations ago. The gene flows from each ancestral population could be zero at a specific generation. POP *k* represents the reference population *k*.

Let *I*(*t*) be the ancestry proportion of the admixed population at generation *t* inherited from the previous generation, and then we get *I*(*t*) as

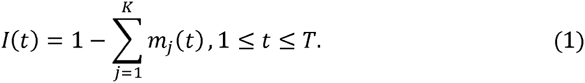

Denote *H*_*i*_(*t*) as the total ancestry proportion of the *ith* ancestral population in the admixed population at *t* generation, then

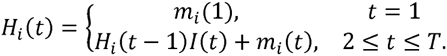

Recursively, we can get

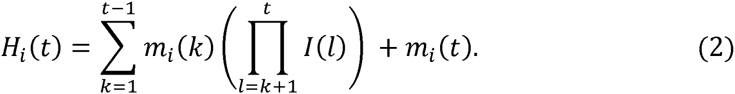

Define *s*_*i*_(*t*) as the survival proportion of the ancestral tracks at generation *T* from the *ith* ancestral population that entered at generation *t*, then

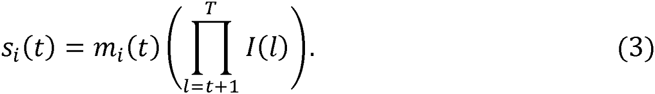

Generally, we assume that the ancestral populations are homogeneous, and recombination among segments from the same ancestry does change the length of the tracks, but it is not “observable” to us, thus the length of tracks seems unchanged, and only these recombination events among different ancestries produce “observable” changes. (Figure S1) Here we explicitly take these “unobservable” changes into consideration and adjust the recombination rate accordingly as following. Define the recombination among tracks from different ancestral populations as effective recombination, let, *u*_*i*_(*t*) be the effective recombination rate for tracks from the *ith* ancestral population that entered at generation *t*, then

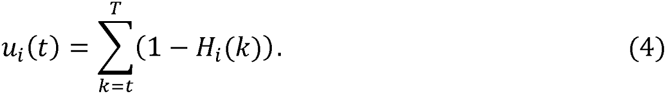

If the end is ignored, a chromosome from the *ith* ancestral population that entered at generation *t* is expected to be split into *u*_*i*_(*t*) pieces per unit length (unit in Morgan). Then the contribution of ancestral tracks from *ith* ancestral population that entered at generation *t* to the admixed population is proportional to *u*_*i*_(*t*)*s*_*i*_(*t*). Let *X*_*i*_ be the length of ancestral tracks of the *ith* ancestral population at generation *T*, and *f*_*i*_(*x*; *M*) be the distribution of *X*_*i*_, then

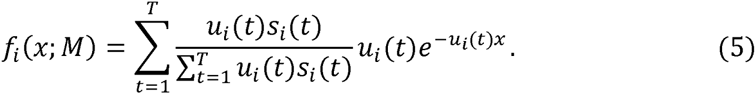

Due to the limited accuracy in local ancestry inference, only those relatively long tracks are reliable (Pool and Nielsen 2009; Gravel 2012). Therefore, we are also interested in the conditional length distribution of ancestral tracks longer than a specific threshold *C*,

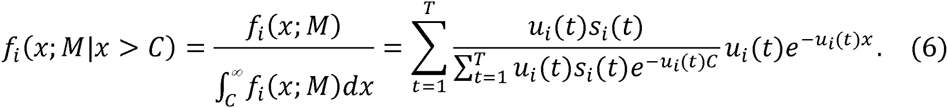

With the length distribution of ancestral tracks (Formula (5)), we can easily deduce the expectation and variance of *X*_*i*_ as

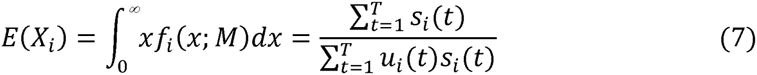

and

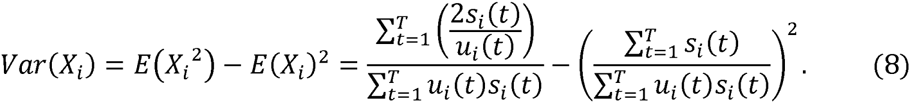

### Parameter Estimation and Model Selection

As the parameters of admixture events are fully determined by the matrix *M*, once *M* was accurately estimated, we could fully recover the history of the admixed population. MLE can be used to obtain the estimation of *M*. By utilizing the ancestral tracks inferred from the data, a likelihood function can be computed with the length distributions of ancestral tracks. The log-likelihood function of ancestral tracks from *ith* ancestral population *L*_*i*_(*M*) has the form

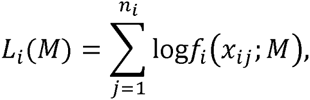

where {*x*_*ij*_}_1≤*j*≤*n*_*i*__ are the observed length of tracks from the *ith* ancestral population in the admixed population. Then the log-likelihood function of ancestral tracks of the admixed population *L*(*M*) is

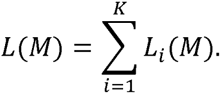

Then the estimator of *M* is

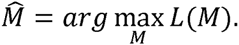

where *M* satisfies the properties in the above subsection.

However, with the increase in the number of parameters, it is complex and time-consuming to find the optimal solution and too many parameters could lead to over-fitting. In a real situation, we can propose several candidate models with prior knowledge in which the number of parameters is dramatically reduced, thus the problem is simplified to estimating parameters for each candidate model and selecting the most suitable one. If we have obtained the parameters of the candidate models, we can compare the models in a pair-wise fashion by using either AIC or LRT. Here, the two models are regarded to be ‘nested’ if one of the models constitutes a special case of the other (Lewis F *et al.* 2011). When the two competing models are nested, we use LRT to select the model; otherwise we use AIC.

## THREE TYPICAL MODELS

### The Distribution of Ancestral Tracks under a HI, GA, and CGF Model

In this subsection, we demonstrate that, with the length distribution of the general model, we can easily deduce the length distributions of ancestral tracks under three typical models aforementioned in previous studies (Jin *et al.* 2012). By restricting the number of ancestral populations to be two in the general model, if only one pulse of gene flow is allowed, the model turns into a HI model; if extra equal gene flows from both ancestral populations are allowed, the model turns into a GA model; if extra equal gene flows from only one of the ancestral populations are allowed, the model turns into a CGF model (Figure S2). Thus these three models are all special cases of our general model. There are only two parameters in each of the three models: the admixture proportion *m* and the admixture time *T*. Easily we can obtain the distribution of the ancestral tracks of these three models from Formula (5). The detailed calculation is in Supporting Information.

For simplicity, we define the ancestral population with the minor ancestry contribution as population 1, and the corresponding proportion is *m*. For the HI model (Figure S2 [A]), the ancestry proportions from population 1 and population 2 at generation *t* are

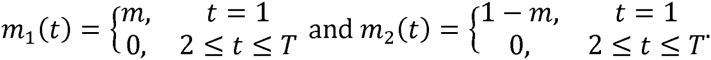

Then the length distribution of ancestral tracks from population 1 is

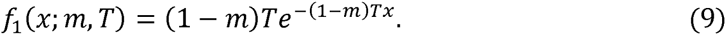

We can also get the expectation and variance of the length of the ancestral tracks from Formula (7) and Formula (8),

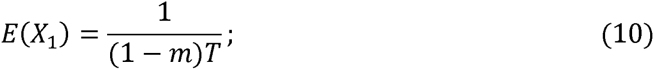

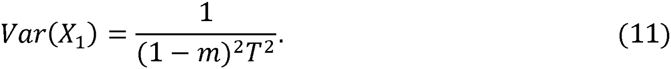

Substituting *m* with 1-*m* in Formulas (9), (10), and (11), we can obtain the length distribution, expectation, and variance of ancestral tracks from population 2, respectively. These two distributions are identical to the ones in previous studies (Gravel 2012; Jin *et al.* 2014).

For the GA model (Figure S2 [B]), the ancestry proportions from population 1 and population 2 at generation *t* are

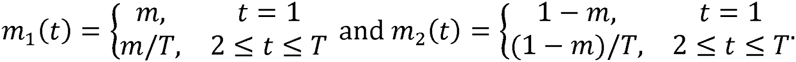

Then the length distribution of ancestral tracks from population 1 is

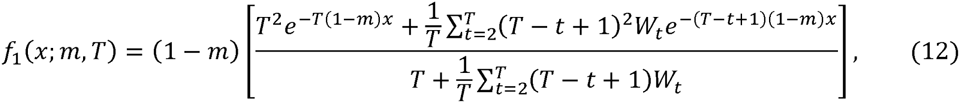

where 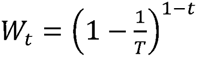. The expectation and variance of the ancestral tracks are

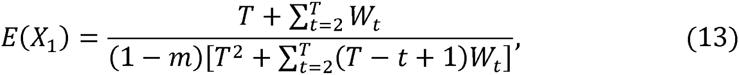

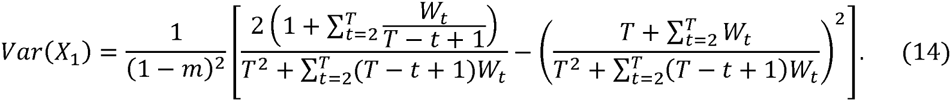

Substituting *m* with 1 − *m* in Formulas (12), (13), and (14), we can get the length distribution, expectation and variance of ancestral tracks from population 2, respectively.

For the CGF model (Figure S2 [C]), the ancestral population that contributes only one pulse of gene flow is treated as a gene flow recipient and the one that contributes continuously as gene flow donor. Here, we divide the CGF model into two sub-models. If population 1 is a gene flow recipient, we denote it as a CGFR model; otherwise we denote it as a CGFD model.

In the case of a CGFR model, the ancestry proportions from population 1 and population 2 at *t* generation are

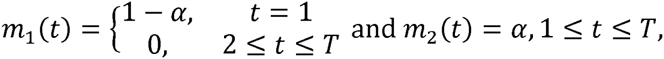

where *α* = 1–*m*^1/*T*^. Then the length distributions of ancestral tracks from the two ancestral populations are

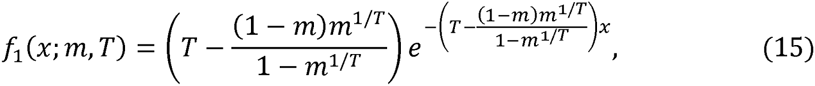

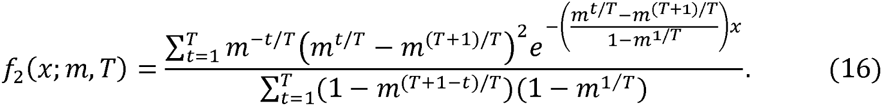

The expectations and variances of the ancestral tracks are

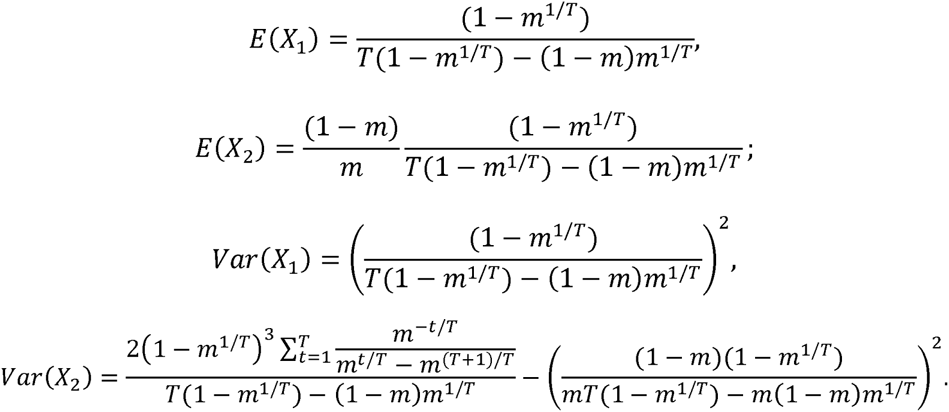

In the case of a CGFD model, we just replace *m* for 1-*m* in Formula (15) and (16), and obtain the length distribution of ancestral tracks from population 2 and population 1, respectively.

### Parameter Estimation and Model Selection under HI, GA, and CGF Models

As discussed above, there are only two parameters *m* and *T* in the HI, GA and CGF models. As for *m*, with the inferred ancestral tracks in an admixed population, we divide the total length of tracks from population 1 by the total length of tracks, and obtain an estimator 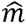 of the admixture proportion. Interestingly, from the expectation of the ancestral tracks from two ancestral populations in the HI, GA and CGF models, we find that the expectation ratio between population 1 and population 2 relies only on *m*,

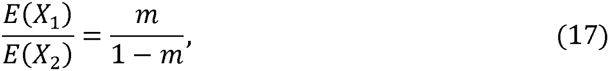

thus we provide an alternative way to obtain the estimator 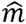,

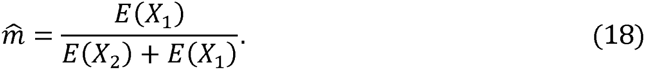

Generally, if there are only two ancestral populations, Formula (17) always holds whatever the admixture model is. The proof is in Supporting Information.

As for admixture time *T*, the estimation relies on the model assumed. Different models give different estimations of *T* so that we first need to assume a model. Here we regard the HI, GA, CGFR and CGFD models as the candidate models, and use MLE to estimate the admixture time *T* and AIC to select the optimal model as following: First, by utilizing the inferred ancestral tracks, we calculate the admixture proportion and determine population 1; secondly, with the estimator 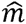, maximizing the likelihood under model assumption HI, GA, CGFR, and CGFD, we obtain the maximum likelihoods *L*_*max*_ (HI), *L*_*max*_ (GA), *L*_*max*_ (CGFR) and *L*_*max*_ (CGFD) and corresponding optimal times 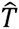. Because each pair of these models is not nested, thus here we use AIC to select the optimal model. The value AIC can be calculated by the formula

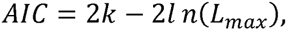

where *k* is the number of parameters and *L*_*max*_ is the maximized value of the likelihood function. The number of parameter of these models are the same, thus at the end of the comparison, we find that the problem is equivalent to finding the model with the highest likelihood. Thus, the model with the highest likelihood is chosen as the optimal model, and the corresponding parameters as the final results. These routines are implemented in our *AdmixInfer*. We also apply the bootstrapping technique in *AdmixInfer* and give a bootstrap estimation and confidence interval (CI) of the admixture time.

## MATERIALS AND METHODS

### General Settings of Simulation Studies

Simulation studies were performed to evaluate the correctness of the length distribution of ancestral tracks under the general model, and the performance of *AdmixInfer* under three typical models. The following settings remained the same under all situations simulated if with no further modification: The population size of the admixed population was simply set to 5,000 and remained constant in our simulations and the length of simulated chromosome was 3.0 Morgan, which approximated the length of chromosome 1 of the human genome. We simulated one chromosome each time and a pair of chromosomes represented an “individual.” At the end of simulation, 400 “individuals” (genome length of an “individual” approximated 1/10 of the length of a human individual) were sampled and the ancestral tracks were directly recorded.

### Evaluate the Theoretical Distribution under the General Model

To test the accuracy of the length distribution of ancestral tracks under the general model, we simulated several general representative cases with three or four ancestral populations and one or more waves of admixture. Simulation 1A was to simulate one pulse admixture with three ancestral populations: admixture started 50 generations ago with proportions 10%, 30% and 60%, respectively. Simulation 1B was to simulate discrete multiple-waves admixture with three ancestral populations: the admixture started 50 generations ago with initial proportions 10%, 40% and 50%, respectively, and population 1 contributed extra 10% ancestries each 10 generations later. Simulation 1C was to simulate continuous multiple-waves admixture with three ancestral populations: the admixture started 50 generations ago with initial proportions 10%, 30% and 60%, respectively; population 1 contributed an extra 0.2% ancestries for each generation afterwards. And simulation 1D was to simulate multiple-waves admixture with four ancestral populations that arrived at different times: populations 1 and 2 firstly admixed 50 generations ago with proportions 40% and 60%, population 3 entered 40 generations ago with proportion 20%, and population 4 entered 30 generations ago with proportion 10%. The simulations were repeated 5 times, and the ancestral tracks in the admixed population were recorded and the length distribution was compared to the theoretical distribution. Detailed parameters for the simulations were provided in Table S1.

### Evaluate the Performance of *AdmixInfer*

Then we focused on evaluating the performance of *AdmixInfer* under three typical models; i.e. HI, GA and CGF. The proportions of admixture varied from 10% to 50% in steps of 10% for the symmetric admixture models (HI and GA) and varied from 10% to 90% in steps of 10% for the asymmetric admixture model (CGF). We set the admixture time as 5, 10, 20, 30,…, 200 generations. The ancestral tracks in admixed populations were also recorded as previous simulations. Each simulation here was repeated 10 times and, in total, 4,200 simulations were carried out under HI, GA and CGF models. *AdmixInfer* was applied to the simulated data with the default settings; the results were recorded and summarized.

In real situations, we could only accurately infer the ancestral tracks longer than a specific threshold due to methods’ limitations in local ancestry inferences. To make our method more feasible to real cases, we also evaluated the robustness of our method under different thresholds ranging from 0 centi-Morgan (cM) to 2 cM in step of 0.1 cM, with the dataset simulated in previous evaluations.

We also evaluated the performance of *AdmixInfer* with different sample sizes. We simulated populations starting with the admixture of 50 and 100 generations ago under HI, GA, and CGF models, with admixture proportions 30%:70%. At the end of the simulation, 10, 20, 50, 100, 200, and 500 “individuals” were sampled, corresponding to 1, 2, 5, 10, 20, and 50 human samples. *AdmixInfer* was applied to the simulated dataset without discarding short tracks.

Finally, we tested the performance of our method with data simulated by real populations and inferred ancestral tracks. Simulations were carried out with real populations YRI and CEU as parental populations under different models (30% YRI ancestry and 70% CEU ancestry) with admixture time 10, 20, 50 and 100 generations. Here we simulated with the data of chromosome 1 and sampled 25 “individuals” at the end of the simulation, and each simulation was repeated 10 times. Then the local ancestry of the simulated populations was inferred by HAPMIX (Price *et al.* 2009). With the derived ancestral tracks, *AdmixInfer* was used to select the optimal model and estimate generations accordingly with the tracks longer than 1 cM.

### Apply to Real Datasets

We applied our method to some real datasets. First, the histories of African Americans and Mexicans are relatively clear, thus they can be used to test the performance of our method. The data of African Americans, Mexicans and reference populations CEU and YRI were obtained from HapMap project phase III (International Hapmap *et al.* 2010), and the reference populations that represented American Indian ancestry were obtained from HGDP dataset. Then we also applied our method to several HGDP populations from South Asia (Li *et al.* 2008), which showed evidence of population admixture from previous studies (Patterson *et al.* 2012; Hellenthal *et al.* 2014). Haplotype phasing was performed by Shapeit 2 (Delaneau *et al.* 2012). Local ancestry was inferred by Hapmix (Price *et al.* 2009). According to the prior knowledge, the generations settings in Hapmix inference were 10, 20 and 50 for African Americans, Mexicans and South Asian populations, respectively (Hellenthal *et al.* 2014). *AdmixInfer* was used to select the optimal model and admixture time accordingly with the tracks longer than 1 cM. We also performed bootstrapping 100 times to obtain confidence of model selection, and calculated the 95% confidence intervals of the generations inferred.

## RESULTS

### Theoretical and Simulated Distributions of Ancestral Tracks Match Well

With the length distributions of ancestral tracks under the general model, we could easily sketch the curves of theoretical length distributions of ancestral tracks under a given model (Figure 2, solid line). By putting the theoretical and simulated length distributions of ancestral tracks together, we clearly observed that the theoretical and simulated distributions of ancestral tracks matched well, for all the situations simulated and all the replicates (Figure 2). It showed that the theoretical length distribution of ancestral tracks for the general model, which we deduced, was reasonable and accurate.

**Figure 2.**
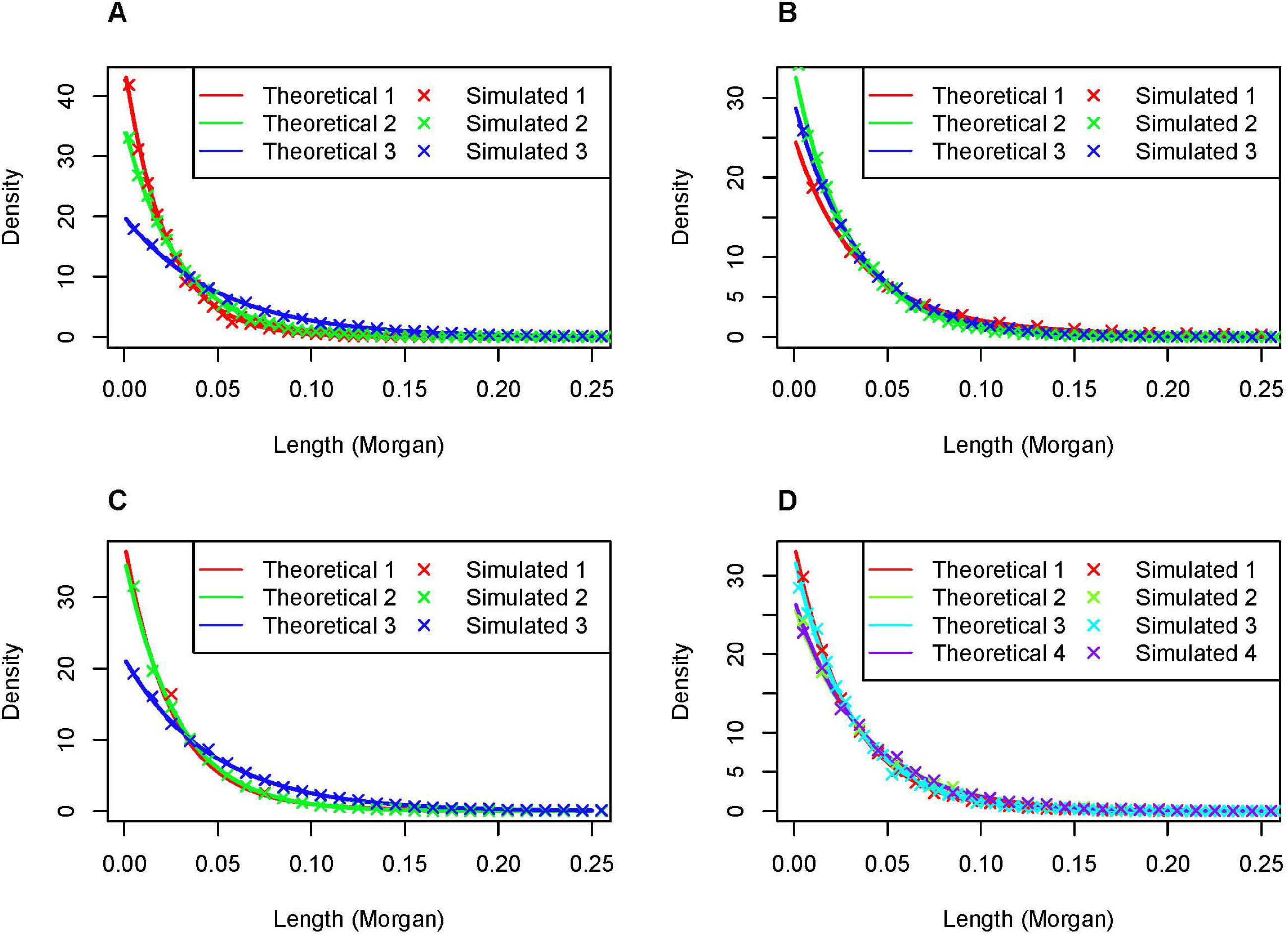
Theoretical and simulated distributions of ancestral tracks under some representative admixture scenarios. A: three reference populations admixed once at 50 generations ago; B: three reference populations admixed at 50 generations ago and population 1 contributed an extra 10% ancestry each 10 generations later; C: three reference populations admixed at 50 generations ago and population 1 contributed an extra 0.2% ancestry every generation later; D: two reference populations admixed 50 generations ago, the third reference population contributed 20% ancestry 40 generations ago, and the fourth reference population contributed 10% ancestry 30 generations ago.

### *AdmixInfer* Performs well in Parameters Estimation and Model Selection

With the extensively simulated data, we could systematically evaluate the performance of our method in parameters estimation and model selection. For the simulated admixed populations under the HI model or CGFD model, our method could always distinguish the right model in all our simulations; for the CGFR model, our method could distinguish the right model with accuracy of 97.0%; and for GA model, our method could distinguish the right model with accuracy of 93.0% (Table 1). Moreover, the specificity of our method was over 97% for all the situations simulated. The sensitivity of the GA and CGFR models and the specificity of the HI and CGFD models under different admixture proportions and different admixture times were shown in Figure S3. We found that the simulations in which our method could not distinguish the right ones were mostly observed in these simulations with very recent admixture times and small admixture proportions (Figure S3 and Table S2-S6). We also found that almost all CGFR models were only wrongly distinguished as HI models, and GA models as CGFD models (Table S6). This is also reasonable, because the CGFR model is close to the HI model, so that CGFR model is more likely to be distinguished as a HI model. The same reason applies for that the GA model being wrongly distinguished as the CGFD model.

**Table 1.** The accuracy of our methods in model selection under three typical models.

| Model | TP | FN | TN | FP | Sensitivity (%) | Specificity (%) |
| --- | --- | --- | --- | --- | --- | --- |
| HI | 1050 | 0 | 3120 | 30 | 100.0 | 99.0 |
| GA | 977 | 73 | 3150 | 0 | 93.0 | 100.0 |
| CGFR | 1019 | 31 | 3150 | 0 | 97.0 | 100.0 |
| CGFD | 1050 | 0 | 3076 | 74 | 100.0 | 97.7 |

HI: Hybrid isolation model; GA: Gradual admixture model; CGFR: Continuous gene flow model (population 1 as recipient); CGFD: Continuous gene flow model (population 1 as donor). TP: True positive; FP: False positive; TN: True negative; FN: False negative. Sensitivity=TP / (TP+FN); Specificity=TN / (TN+FP).

Note that there were only two parameters *m* and *T* for the three typical models. Our method also performed well in estimating parameters *m* and *T* for the three typical models. Regarding the admixture proportion m, estimations were very close to the pre-settings in simulations, and the small deviation was due to random drift in simulation with finite population sizes and only a subset of individuals being sampled at the end of the simulation. Time estimations for wrongly distinguished models were meaningless, and thus should be discarded. Results showed that our method can estimate admixture times with high accuracy (Figure 3, Figure S4-S7, and Table S2-S5). For HI and CGFD models, results showed high consistency with the time simulated, while a slight underestimation occurred for GA and CGFR models. For all these models that were simulated, the time estimated for a small proportion was less accurate than that for a larger proportion. We defined the relative errors between true admixture times and inferred admixture times as

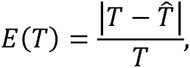

**Figure 3.**
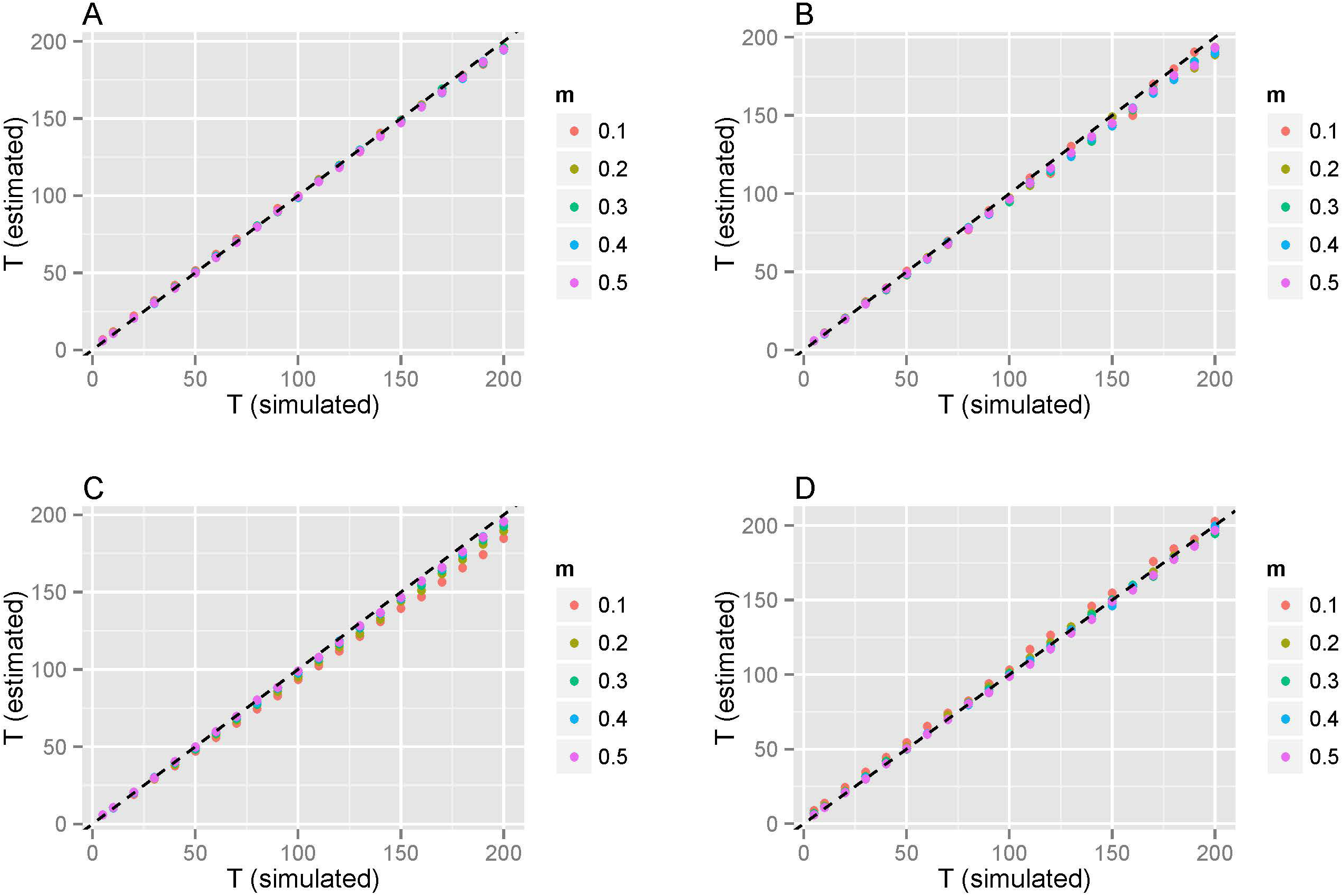
Mean generations estimated from simulation. Each dot denotes the mean generation of ten simulation replicates. A: mean generations estimated under the HI model; B: mean generations estimated under the GA model; C: mean generations estimated under the CGF model (population 1 as gene flow recipient); and D: mean generations estimated under the CGF model (population 1 as gene flow donor). Different colors represent different simulated proportions of population one.

where *T* is the true admixture time and *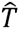* is the estimation of the admixture time. Under the situation of a certain admixture proportion and a certain model, we defined the average relative error 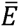 on different values of admixture time as

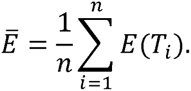

We found that when the admixture proportion was 0.1, the relative errors of CGFR and CGFD were 6.43% and 5.89%, respectively. For the other cases, the relative errors were all less than 4% (Table 2). In conclusion, no matter the model selection or parameters estimation, our method performed well.

**Table 2.**
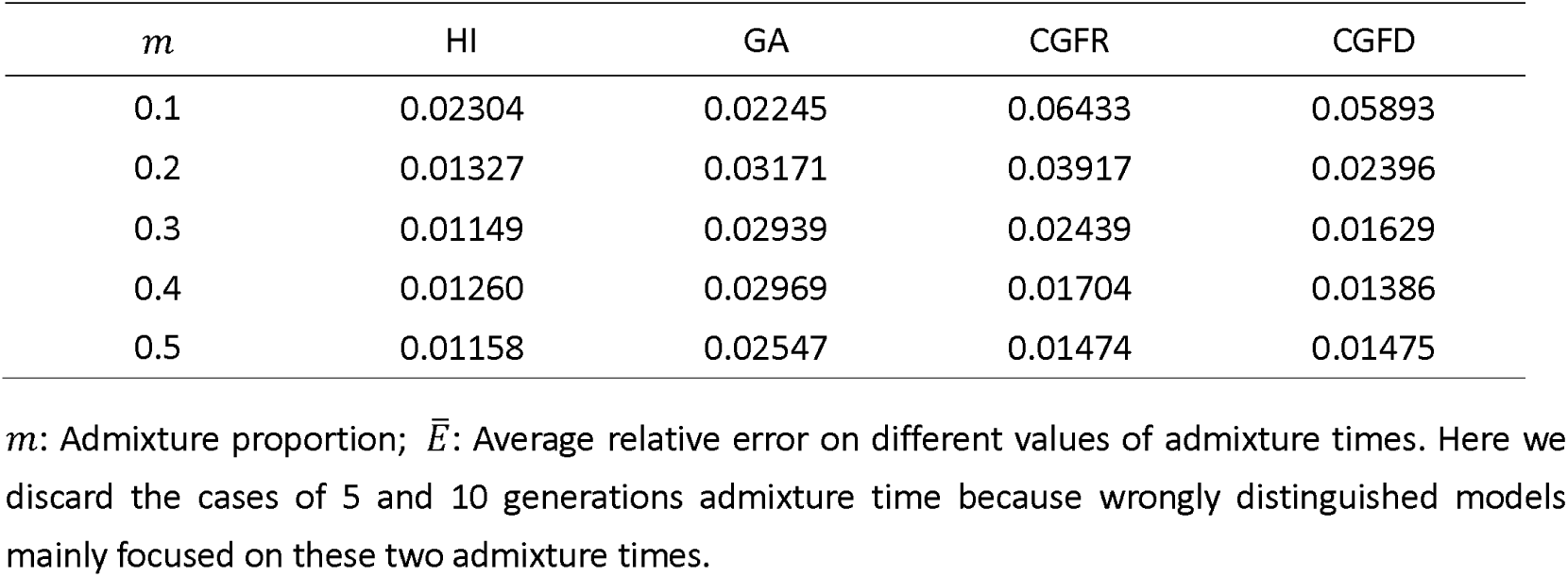
The average relative error 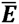 for different values of admixture proportions.

### Robustness for Different Thresholds

To test the robustness of our method for different thresholds, we tested our method under different thresholds varying from 0 cM to 2 cM in steps of 0.1 cM. The results showed that our method was robust to thresholds, except the GA model with a larger time (Figure 4). When a larger threshold is taken, less information is kept for ancient admixture events. Although keeping all the information to estimate admixture times is better, we must balance the trade-off between information and accuracy, because the accuracy of local ancestry inference is not so good for short ancestral tracks due to method limitations. Take HI for an example, the probability *p* of ancestral tracks larger than a specific threshold *C* is

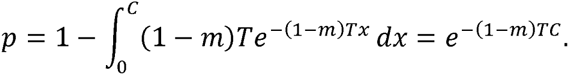

**Figure 4.**
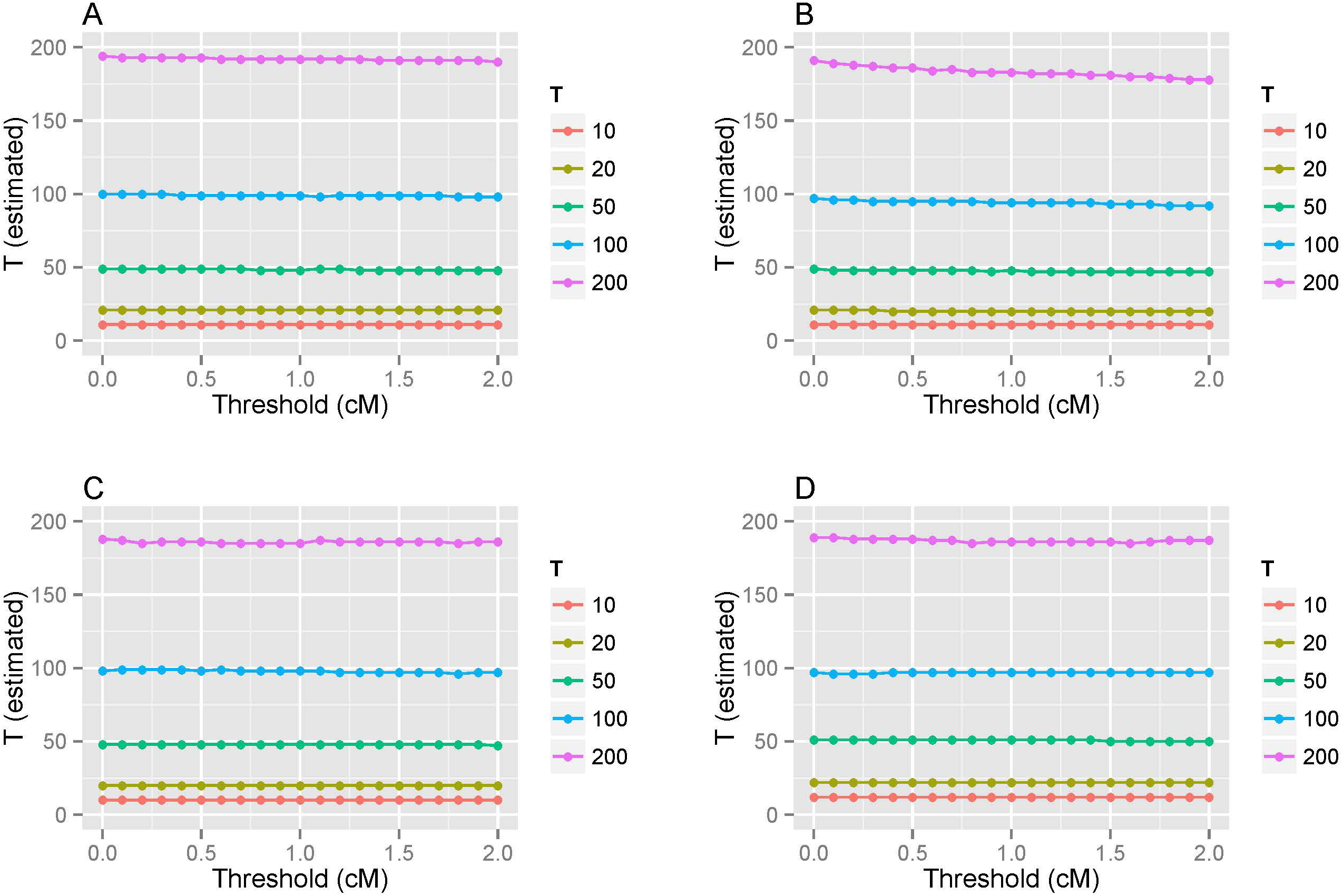
Generation estimated with different thresholds from simulation. Models simulated were HI(A), GA(B), CGFR(C) and CGFD(D). The simulated admixture proportion was 30%. Different colors represent different simulated generations.

Therefore, with an increase in threshold *C*, less information is kept. Here, we provided a reference table of the probability *p* under different admixture times and proportions (Table S7). For example, when *T* = 200, *m* = 0.1, if we set the threshold *C* as 2cM, the probability that the tracks exceed *C* is only 2.7%.

### Small Sample Sizes also Give Good Estimations

To test the performance of *AdmixInfer* with different sample sizes, we evaluated the models with 10, 20, 50, 100, 200, and 500 “individuals” (corresponding to 1, 2, 5, 10, 20, and 50 human samples). Results showed that *AdmixInfer* was insensitive to sample sizes. Even with only one human sample, it could distinguish the right model and estimate the admixture time with high accuracy (Figure 5). However, considering the accuracy of local ancestry, short tracks were usually discarded. The information kept for extremely small sample sizes might not be sufficient to give a clear picture of the history of a population. Therefore, relatively larger sample sizes were recommended.

**Figure 5.**
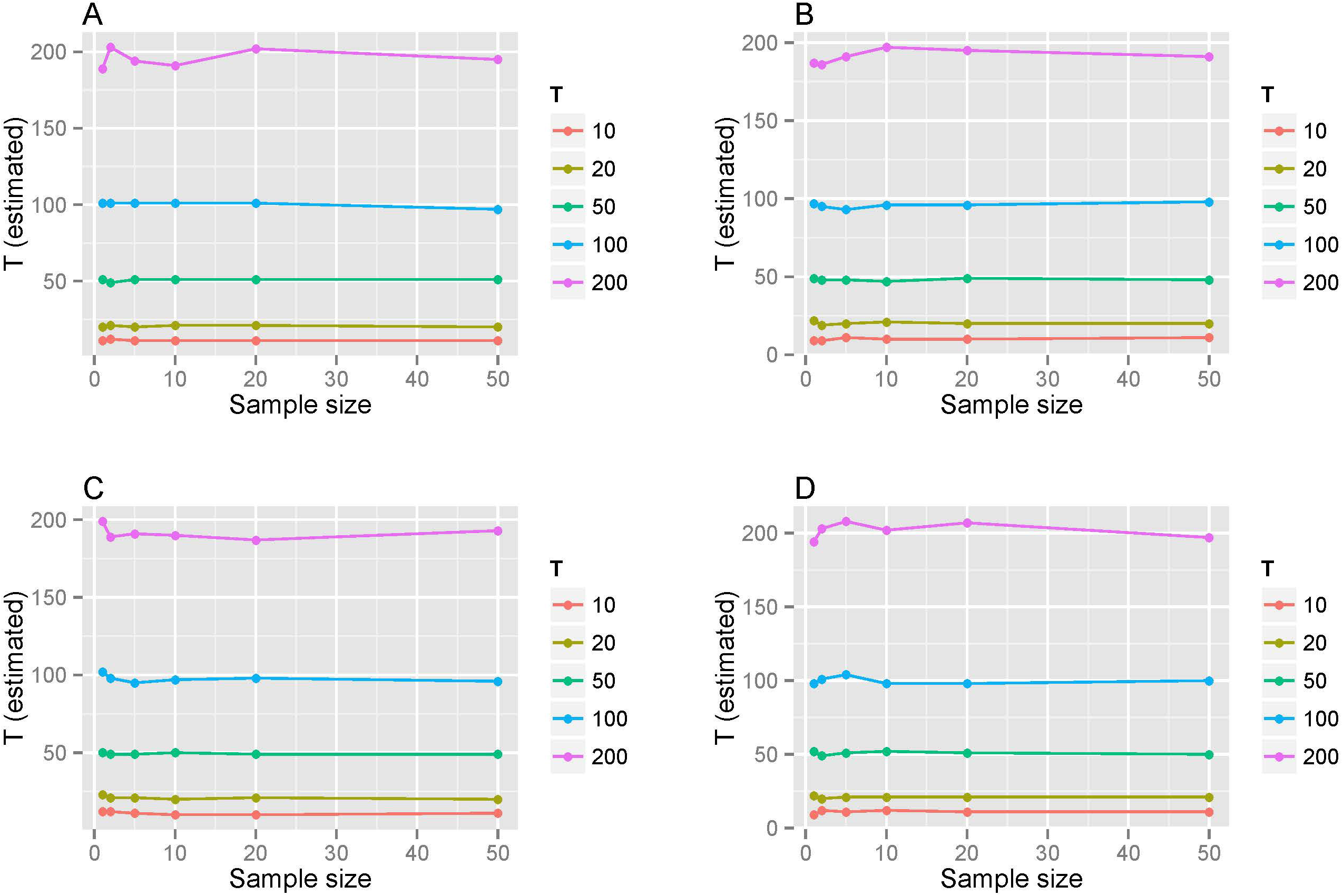
Admixture time in generations estimated with different sample size. Models simulated are HI (A), GA (B), CGFR (C), and CGFD (D). Different colors represent different simulated generations. The simulated admixture proportions was 30%.

### Error Analysis

When we use our method to infer the history of a real admixed population, there are two kinds of errors that may influence the accuracy of inference. The first kind of error is caused by the assumptions in deducing the length distribution of ancestral tracks. In the derivation, for simplicity, we ignored the end of the chromosome and the drift. For this kind of error, we have used the simulation data to demonstrate that the accuracy of the inference was neglectable (Figure 3 and Table 1). The second kind of error is caused by the local ancestry inference. In our study, local ancestral tracks are inferred by HAPMIX software. Here we used the simulation data with ancestry populations YRI and CEU to analyze the influence of this kind of error. For all the cases we simulated, only the HI model with an admixture time 100 generations was wrongly distinguished as a GA model (Table S8). We also found that for the case of large admixture times, the error of local ancestry inference will cause underestimation of admixture time. When the admixture time is large, the ancestral tracks will be short. However, the method of inferring ancestral tracks cannot effectively determine short tracks. Thus, it will influence the accuracy of our method in inferring admixture times and model selections.

### Real Data Analysis

Our method for parameters estimation and model selection under three typical models implemented in *AdmixInfer* was applied to a real dataset. First, African American was inferred as a GA model and the admixture time was 12 generations ago (Table 3). When 29 years per generation was assumed according to previous investigation (Fenner 2005), it was about 350 years ago, which was consistent with the recorded history that most African ancestors arrived in America (via slave trading) after the seventeenth century. The slave trade continued until the nineteenth century and after that, many African people settled down in America. Gene flows from Africa and Europe would have continuously contributed to the African American gene pool and thus the GA model matched the recorded history well. Similarly, the model for Mexicans was inferred as a GA model and the admixture time was 18 generations ago (522 years before present), which was also consistent with the time of the exploration of the new world and the arrival of Europeans. The GA model indicates continuous contact and admixture of Europeans and American Indians.

**Table 3.** Admixture time and model inferred for real datasets.

| POP1 | POP2 | Admixed pop | $m$ | Model | $T$ | 95%CI |
| --- | --- | --- | --- | --- | --- | --- |
| CEU | YRI | ASW | 0.246748 | GA (99%) | 12 | [12,12] |
| CEU | AMI | MEX | 0.477949 | GA (100%) | 18.02 | [17.99, 18.05] |
| Japanese | Sardinian | Uygur | 0.569544 | GA (100%) | 70.43 | [70.2, 70.66] |
| Japanese | Sardinian | Hazara | 0.558653 | GA (100%) | 67.16 | [67.01, 67.31] |
| Japanese | Sardinian | Balochi | 0.197582 | CGF (100%) (Sardinian as donor) | 104.7 | [104.5, 104.8] |
| Japanese | Sardinian | Brahui | 0.189526 | CGF (100%) (Sardinian as donor) | 105.2 | [105, 105.3] |
| Japanese | Sardinian | Burusho | 0.339483 | CGF (100%) (Sardinian as donor) | 96.52 | [96.34, 96.7] |
| Japanese | Sardinian | Kalash | 0.173549 | CGF (100%) (Sardinian as donor) | 100.5 | [99.27, 101.6] |
| Japanese | Sardinian | Pathan | 0.257682 | CGF (100%) (Sardinian as donor) | 98.49 | [98.28, 98.7] |
| Japanese | Sardinian | Sindhi | 0.271986 | CGF (100%) (Sardinian as donor) | 107.5 | [107.4, 107.7] |

POP1: Reference population one; POP2: Reference population two; Admixed pop: Admixed population; $m$ : Admixture proportion of reference population one; Model: Inferred admixture model, percentage in the parenthesis is the support rate in 100 times bootstrapping; $T$ : estimated admixture time in generation; 95%CI: 95% confidence interval of the estimated admixture time; AMI: Combined dataset of populations Maya and Pima which represent American Indian ancestry.

Finally, we studied the admixture histories of several HGDP populations from South Asia. Previous studies have shown that the populations from South Asia have admixed ancestries mainly from Europe and East Asia (Li *et al.* 2008; Hellenthal *et al.* 2014). Regarding the admixture proportions, our estimations based on *AdmixInfer* (Table 3) were consistent with previous estimations from the three-population test (Patterson *et al.* 2012). Regarding the admixture model and time, the populations Balochi, Brahui, Burusho, Kalash, Pathan and Sindhi from Pakistan in South Asia were all inferred to as CGF (Sardinian as donor), which indicated extra gene flows from European ancestry after initial admixture. The initial admixture times estimated ranged from 107 to 96 generations ago. When 29 years per generation was assumed according to previous investigations (Fenner 2005), these estimations (1103BC-784BC) coincided with the migration of Indo-Aryan speaking people into the Indian subcontinent. Extra gene flows from European ancestry might be contributed to by the rising of empires during the following centuries. In the case of the populations of Hazara and Uygur, they not only showed very similar admixture proportions, but also showed the same admixture model and very close admixture times; i.e., 67 and 70 generations ago, respectively. The Hazara population mainly settled in Afghanistan and Pakistan, while the Uygur population mainly settled in West China, and both populations are connected along the Silk Road. It was feasible to receive continuous gene flows from both European and East Asian ancestries. These similarities also indicated a possible close relationship or shared histories between these two populations.

In summary, our method showed good performance in inferring the admixture history of African Americans and Mexicans. The admixture scenarios in South Asian populations were more complex than expected. However, with our method, the analysis could shed light on the mysterious histories of these populations.

## DISCUSSION

In this work, we proposed a general model to describe the admixture history with multiple ancestral source populations and multiple-wave admixtures. We showed the length distribution of ancestral tracks and some of its useful properties under this general model. We thus provided a theoretical framework to study population admixture history. With the general framework, we focused on studying three special cases of the admixture models (HI, GA, and CGF) and developed a method to estimate the admixture proportion, admixture time and determine the optimal model simultaneously. Our simulations showed that the theoretical distribution of ancestral tracks was consistent with our theoretical prediction, and our method was precise and efficient in inferring population history under three typical models.

In the efforts of model selection, we found that the simulations in which our method was not able to determine the correct model, were mostly those cases with recent admixture times and minor admixture proportions. The possible reason for incorrect determination was that we ignored the chromosome ends in deducing the theoretical length distribution. When the admixture proportion and times were small, the chromosomes without “observable” recombination were over-represented in the ancestral tracks. (Figure S8). Our further simulations showed that when the chromosome length increased, the accuracy of our method was enhanced. Furthermore, we note that the length distributions of ancestral tracks have no relationship with the population size. Thus, the change of population size does not affect the time estimation. Simulations under different demographic models also supported it (Figure S9).

The efficiency of our method could also be influenced by the validity of the local ancestry inference. To improve the reliability of the inference, we suggest using the ancestral tracks longer than a certain threshold *C*. However, when the threshold became large, some ancient admixture information disappeared rapidly. In principle, if short ancestral tracks could be precisely detected, our method is promising in recovering even more ancient admixture history, such as the admixture between modern human and Neanderthals (PrÜfer *et al.* 2014; Sankararaman *et al.* 2014).

Though we proposed a general framework and relevant principles to infer the population history under the general model, finding optimal estimation for parameters is a challenging work with high dimensionality. Currently, our method implemented in *AdmixInfer* is focusing on the three typical models. For the real admixed populations, the admixture history is always complex, such as discrete multiple-waves admixture. Under such circumstances, the length distribution of ancestral tracks under the general model is still broadly useful and applicable. Therefore, based on this framework, to infer more complicated admixture history is a problem to be solved in the future.

## ACKNOWLEDGEMENTS

This work was supported by the Strategic Priority Research Program of the Chinese Academy of Sciences (CAS) (XDB13040100), by National Natural Science Foundation of China (NSFC) grants (91331204 and 31171218), 973 Project (2011CB808000), the Fundamental Research Funds for the Central Universities (2011JBZ019), the National Excellent Doctoral Dissertation Foundation of PR China (FANEDD 201312), National Center for Mathematics and Interdisciplinary Sciences of CAS, and the Key Laboratory of Random Complex Structures and Data Science, CAS (2008DP173182). S.X. also gratefully acknowledges the support of the National Program for Top-notch Young Innovative Talents of The “*Wanren Jihua*” Project.

## References

Akaike, H., 1998 Information theory and an extension of the maximum likelihood principle, pp. 199–213 in Selected Papers of Hirotugu Akaike. Springer.

Delaneau, O., J. Marchini and J. F. Zagury, 2012 A linear complexity phasing methodfor thousands of genomes. Nat Methods 9: 179–181.

Fenner, J. N., 2005 Cross-cultural estimation of the human generation interval for use in genetics-based population divergence studies. Am J Phys Anthropol 128:

Gravel, S., 2012 Population genetics models of local ancestry. Genetics 191: 607–619.

Hellenthal, G., G. B. Busby, G. Band, J. F. Wilson, C. Capelli et al., 2014 A genetic atlas of human admixture history. Science 343: 747–751.

International Hapmap, C., D. M. Altshuler, R. A. Gibbs, L. Peltonen, D. M. Altshuler *etal*., 2010 Integrating common and rare genetic variation in diverse human populations. Nature 467: 52-58.

Jin, W., R. Li, Y. Zhou and S. Xu, 2014 Distribution of ancestral chromosomal segments in admixed genomes and its implications for inferring population history and admixture mapping. Eur J Hum Genet 22: 930–937.

Jin, W., S. Wang, H. Wang, L. Jin and S. Xu, 2012 Exploring population admixture dynamics via empirical and simulated genome-wide distribution of ancestral chromosomal segments. Am J Hum Genet 91: 849–862.

Lewis F, Butler A and G. L, 2011 A unified approach to model selection using the likelihood ratio test. Methods in Ecology and Evolution 2: 155–162.

Li, J. Z., D. M. Absher, H. Tang, A. M. Southwick, A. M. Casto et al., 2008 Worldwide human relationships inferred from genome-wide patterns of variation. Science 319: 1100–1104.

Loh, P. R., M. Lipson, N. Patterson, P. Moorjani, J. K. Pickrell et al., 2013 Inferring admixture histories of human populations using linkage disequilibrium. Genetics 193: 1233-1254.

Patterson, N., P. Moorjani, Y. Luo, S. Mallick, N. Rohland et al., 2012 Ancient admixture in human history. Genetics 192: 1065–1093.

Pickrell, J. K., N. Patterson, P. R. Loh, M. Lipson, B. Berger et al., 2014 Ancient west Eurasian ancestry in southern and eastern Africa. Proc Natl Acad Sci U S A 111: 2632-2637.

Pool, J. E., and R. Nielsen, 2009 Inference of historical changes in migration rate from the lengths of migrant tracts. Genetics 181: 711–719.

Prüfer, K., F. Racimo, N. Patterson, F. Jay, S. Sankararaman et al., 2014 The complete genome sequence of a Neanderthal from the Altai Mountains. Nature 505: 43-49.

Price, A. L., A. Tandon, N. Patterson, K. C. Barnes, N. Rafaels et al., 2009 Sensitive detection of chromosomal segments of distinct ancestry in admixed populations. PLoS Genet 5: e1000519.

Pugach, I., R. Matveyev, A. Wollstein, M. Kayser and M. Stoneking, 2011 Dating the age of admixture via wavelet transform analysis of genome-wide data. Genome Biol 12: R19.

Sankararaman, S., S. Mallick, M. Dannemann, K. Prufer, J. Kelso et al., 2014 The genomic landscape of Neanderthal ancestry in present-day humans. Nature 507: 354-357.

Wilks, S. S., 1938 The large-sample distribution of the likelihood ratio for testing composite hypotheses. The Annals of Mathematical Statistics 9: 60–62.

Xu, S., W. Huang, J. Qian and L. Jin, 2008 Analysis of genomic admixture in Uyghur and its implication in mapping strategy. Am J Hum Genet 82: 883–894.

